# Factorization-based Imputation of Expression in Single-cell Transcriptomic Analysis (FIESTA) recovers Gene-Cell-State relationships

**DOI:** 10.1101/2021.04.29.441691

**Authors:** Elnaz Mirzaei Mehrabad, Aditya Bhaskara, Benjamin T. Spike

## Abstract

Single cell RNA sequencing (scRNA-seq) is a gene expression profiling technique that is presently revolutionizing the study of complex cellular systems in the biological sciences. Existing scRNA-seq methods suffer from sub-optimal target recovery leading to inaccurate measurements including many false negatives. The resulting ‘zero-inflated’ data may confound data interpretation and visualization. Since cells have coherent phenotypes defined by conserved molecular circuitries (i.e. multiple gene products working together) and since similar cells utilize similar circuits, information about each expression value or ‘node’ in a multi-cell, multi-gene scRNA-seq data set is expected to also be predictable from other nodes in the data set. Based on this logic, several approaches have been proposed to impute missing values in a data set by extracting information from its non-zero measurements. In this study, we apply non-negative matrix factorization to a selection of published scRNA-seq data sets followed by multiplication of the factor matrices to generate idealized ‘completed’ model versions of the data. From the model matrices, we recommend new values where original measurements are likely to be inaccurate and where ‘zero’ measurements are predicted to be false negatives. The resulting imputed data model predicts novel type markers and expression patterns that match orthogonal measurements and field literature better than those obtained from pre-imputation data or alternative imputation strategies.

**Availability and implementation:** FIESTA is written in R and is available at https://github.com/elnazmirzaei/FIESTA and https://github.com/TheSpikeLab/FIESTA.

**Author summary:** In this work, we develop FIESTA, a novel, unsupervised, mathematical approach to impute missing values in scRNA-seq data. For each dataset, we use parts-based, non-negative matrix factorization to break the cells-by-genes expression matrix into optimized component matrices and then multiply these component matrices to generate an idealized, ‘completed’ matrix. The completed matrix has many of the null values filled in because the optimized low rank factors from which it is generated, take multiple cells into account when estimating a particular component, including some cells with positive expression values for genes which are false negatives in other related cells. We also implement scaling and thresholding approaches based on intrinsic data topology for improved interpretability and graphical representation. Overall, FIESTA performs favorably relative to alternative imputation approaches and uncovers gene-gene and gene-cell relationships that are occluded in the raw data. The FIESTA computational pipeline is freely available for download and use by other researchers analyzing scRNA-seq data or other sparse data sets.

## 1 Introduction

Single cell RNA sequencing (scRNA-seq) is a powerful laboratory technique aimed at quantifying the abundance of all the transcripts within individual cells. Although it is now a widely used approach for the identification of cell types and cell states based on characteristic gene expression patterns, scRNA-seq typically suffers from incomplete recovery of the cellular RNA pool within each cell (1-3). Recently, several data imputation approaches have been proposed to address inaccuracy and ‘zero-inflation’ resulting from this transcript dropout effect (2-6). Common to these computational approaches is the idea that missing values can be inferred and corrected by borrowing information from non-zero measurements obtained from similar cells and/or correlated genes. For example, the scImpute approach identifies similar cells by spectral clustering and then assigns a probability that a given zero value represents a dropout event, subsequently recommending a replacement value based on the bimodality and variance of expression distributions in closely clustering cells (4). Another approach, MAGIC, identifies similar cells using an adaptive Markov model in PCA space and subsequently imputes values for each gene using a diffusion model and pre-PCA values in ‘neighboring’ cells (6). The SAVER approach uses uses a bayesian model based on UMI-counts and LASSO penalized regression to predict appropriate replacement values for all genes (2). Another example, DeepImpute, employs a deep neural network machine learning model to identify predictive gene subsets/networks for missing value prediction/imputation(5). A number of other approaches have also been reported (7-11). Together, these many attempts at accurately imputing missing data by borrowing information from the coherency of cell types and gene circuits attest to the widespread interest among sequencing users in the potential to computationally impute missing data that is biologically meaningful.

We recognized that recovering missing values in single cell expression data can be viewed as a sparse matrix completion problem and hypothesized that a recommender system based on non-negative matrix factorization (NMF) could provide an especially effective means to recover missing values since the approach has been widely and effectively used to make missing-value predictions from sparse data matrices in other fields (12-14). Furthermore, NMF has been shown by several groups including our own to be particularly effective in delineating biologically relevant cell types and meaningful cell type-associated gene expression profiles from cell expression data (15-18). It thus presents an attractive approach to factorization-based imputation that is likely to draw from relevant biological substructures in the data. The basic process of matrix completion with NMF (or with Singular Value Decomposition (SVD), as employed by two other recently published approaches, CMF-impute (7) and ALRA (3)) is related to the mathematical procedure by which a matrix can be represented as the product of two factor matrices similar to the way a non-prime number can be represented as the product of its factors. If perfect matrix factors can be found, their product will reproduce the original matrix. However, finding a perfect factorization solution for an empirical gene expression matrix is a potentially intractable process (19), and machine learning techniques are therefore often used to generate optimized approximate factors instead. In the case of a matrix factored by NMF, the optimized factors are two strictly positive component matrices (U and V) where rows in U and columns in V are equal to the height and width dimensions of the input matrix, respectively, and their opposing dimension in each case is equal to a selected rank, k. k can be chosen manually or computationally, optimized to balance dimensionality reduction (lower rank) with resolution (higher rank) (Figure 1).

**Figure 1.**
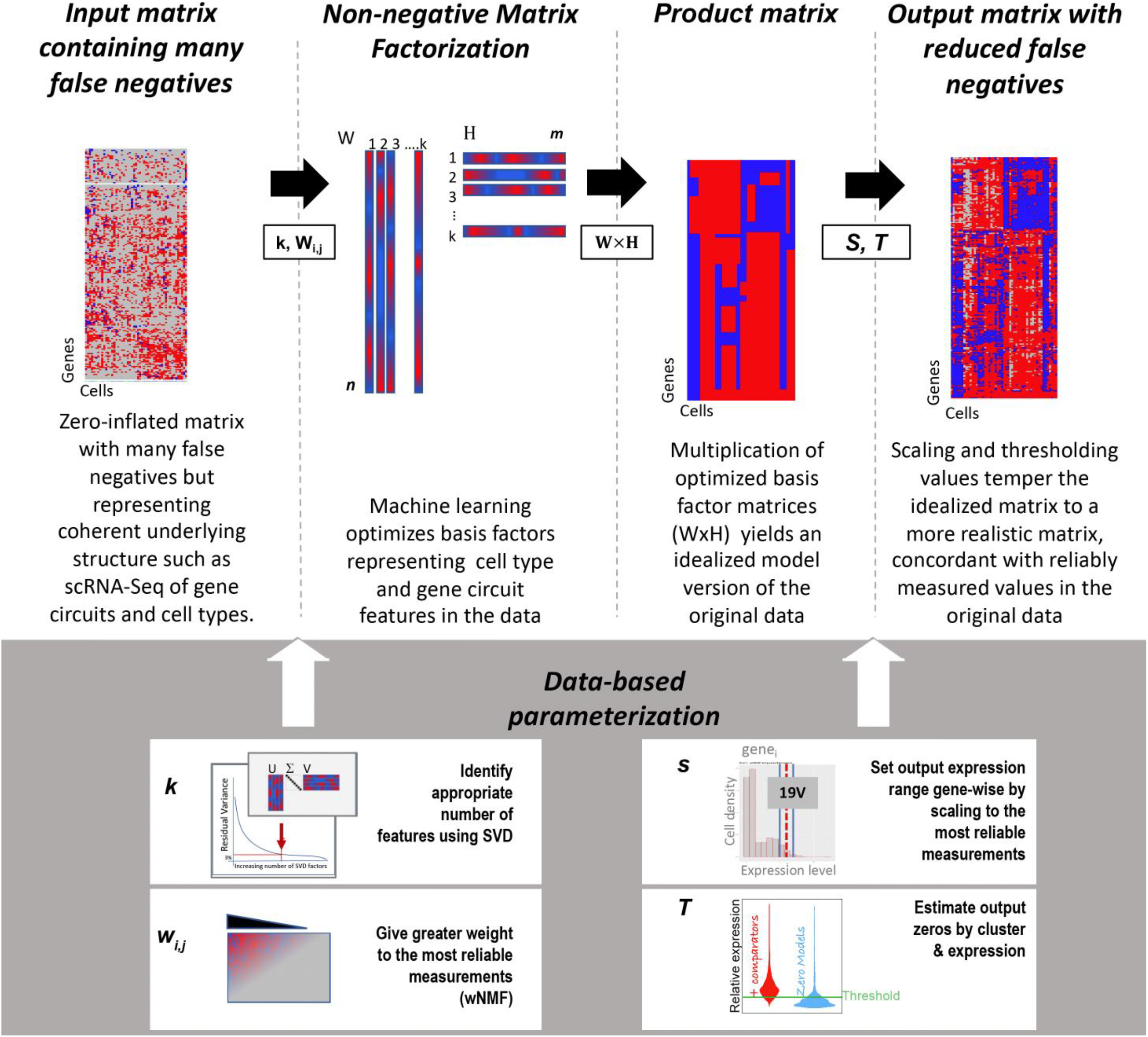
Overview of FIESTA. FIESTA derives scaling and factorization parameters from an input matrix, then factors the matrix with NMF, derives a factor product matrix and applies scaling and thresholding to create a tempered, imputed matrix.

The factors (columns) in U represent generalized gene expression data features that may be expressed to different degrees in different cells; however, as different cells are unlikely to exhibit false nulls at the same individual genes or nodes, the generalizing, optimized factors representing these data (and ultimately their product matrices) tend to fill in any values that were missing in only one or a few cells that share the feature. Thus, the ‘completed’ product matrix serves as an idealization of the original matrix and can be used to recommend corrected values. Here, we present an unsupervised computational pipeline for Factorization-based Imputation of Expression in Single-cell Transcriptomic Analysis (FIESTA) – an NMF-based, machine-learning recommender system for imputation of missing values in scRNA-seq data. FIESTA is based on reconstitutional matrix completion following either of two modified NMF approaches: sparse-NMF (sNMF) (20) or Weighted NMF (WNMF) (21, 22). FIESTA employs factorization rank, gene and cell-weight, and scaling parameters derived from non-zero values in the original normalized matrix and includes an optional thresholding procedure to distinguish false negatives and post involution false positives from likely true negatives. We applied FIESTA to a selection of published data sets and compared its effectiveness to several other published methods. FIESTA performed favorably relative to each of these techniques in recovering expression values known from orthogonal analysis of transcript levels or from literature. FIESTA is also effective across a broad range of initial detection levels and facilitates the resolution of novel cell type markers.

## 2 Materials and methods

### 2.1 Overview and Input data

#### 2.1.1 Data Sets and Format

FIESTA takes annotated single cell expression data in the form of an m*n matrix as input, where m represents the number of genes (rows) and n represents the number of cells (columns) in a given matrix, R. FIESTA then processes this matrix through a factorization and completion algorithm to produce an imputed matrix ImputedR (U_m*k_*(V_n*k_)^T^) (Figure 1). In this study, we used 4 different published data sets to assess the effectiveness of FIESTA, each bearing reasonable expectations of ‘ground-truth’ expression values from orthogonal measurements and published system knowledge. The data sets employed are:

1. a melanoma dropseq data set (12241 genes * 8498 cells) with paired orthogonal quantification of transcripts using in situ hybridization (23),
2. a data set (17059 genes * 6807 cells) from a published study of mouse lung adeno-carcinoma cells involving experimentally manipulated genetics and molecular therapeutic treatments, and bearing associated transcriptional cell state changes (24),
3. a subset (22184 genes * 3562 cells) of our previously published adult mouse mammary epithelial cell scRNA-seq data set, where we have knowledge of tissue specific ‘ground truths’ from our own studies and from the literature (17),
4. Data from isolated peripheral blood mono-nuclear cells (PBMC) (13,458 genes *7,667 cells)(25) as collated by Linderman et al. (3), consisting of well characterized B cells, Monocytes, and T cells.

#### 2.1.2 Pipeline overview

As depicted in Figure 1, FIESTA uses NMF to render input data into a machine-learning-optimized, low-rank representation comprised of basis matrices, U and V. The product of U and V generates an idealized version of the original data. To achieve this FIESTA first estimates overall data complexity (i.e. estimation of suitable k), factorizes the data with NMF and then produces the product matrix (imputed data). Fiesta then identifies critical parameters from the input data in combination with the imputed data for scaling and thresholding, including gene wise expression scales, and cluster-based (i.e., cell type aware) identification of multimodality and likely-true zeros. Thus, FIESTA generates an imputed data set with reasonably modeled expression values and greatly reduced false-negative entries.

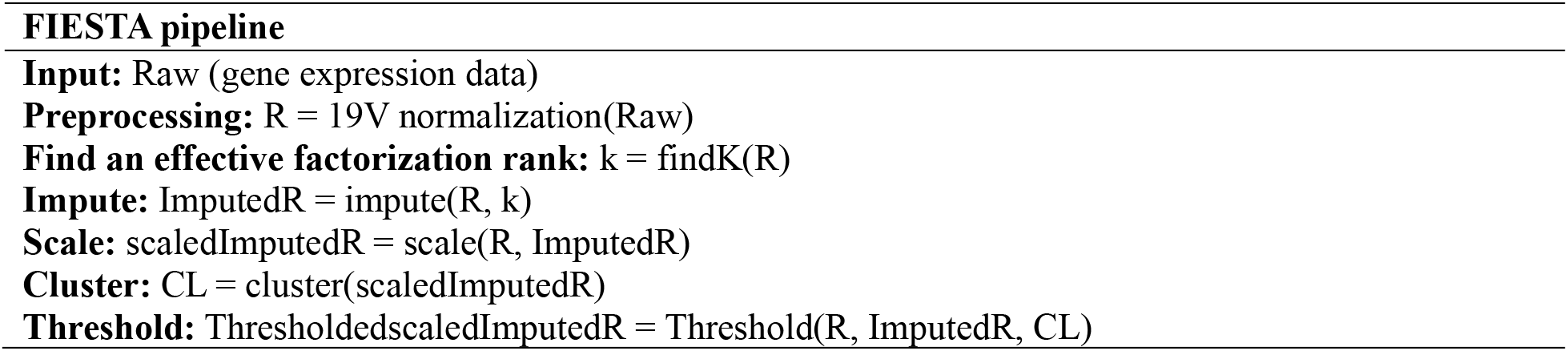

#### 2.1.3 Data pre-processing

Tabulated raw data is first processed by ‘19v’ normalization as in Giraddi et al. (17), a scaling/down-sampling step that normalizes cell-wise sequencing depth without assuming equal overall transcript number in different cells. This processing step takes the 19th ventile of expressed values in each cell (i.e. the sum of the 90th-95th quantile values) as a standard value and derives coefficients for each cell such that multiplying all values in a cell by its coefficient normalizes its 19v to a common set point across all cells. Initially, each element undergoes division by its corresponding 19th ventile value. Cells exhibiting a 19th ventile value less than 10 are excluded. The final coefficient is used to scale all elements uniformly. The computation of this coefficient involves detecting and removing additional outlier cells by applying a z-score threshold of 2 to the 19th ventiles. Subsequently, the 25th percentile of the original 19th ventile value set is chosen as the set point value. Following this, all values are subjected to multiplication by this setpoint divided by their sample 19v coefficient, thereby completing the normalization process. 19v processing is conducted on unlogged data and we recommend logging the data after the imputation step (i.e. *Log*10(*raw* + 1)) for further analysis.

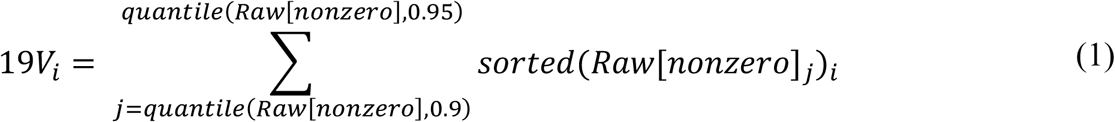

Genes with no positive values in any cell set are withheld from factorization as unimputable due to insufficient information.

### 2.2 Parameter derivation/application and Imputation

#### 2.2.1 Finding an effective factorization rank (k)

FIESTA first recommends a suitable rank (k) by approximating data set complexity using an SVD diagonal matrix and associated variance capture (vc). Although SVD is a factorization approach that is conceptually distinct from NMF and does not have a non-negative constraint, it has been shown to characterize overall gene expression data set complexity to a similar degree as NMF when using similar ranks (26, 27). Nevertheless, the associated k value can be manually adjusted by users for greater or less resolution as desired. As a default setting, FIESTA selects a rank (k) from the diagonal matrix of SVD using the lowest rank k for which >97% of the variance in the data is captured. Thus, FIESTA selects k for each new input matrix R as follows:

Consider D is the diagonal matrix from SVD of the input matrix, and n is the number of cells, then,

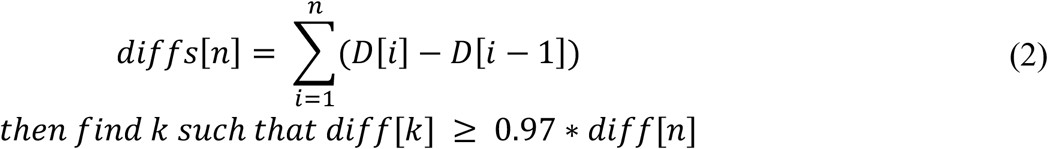

#### 2.2.2 Imputation step

Either of two NMF factorization implementations (WNMF or sNMF) are used to factor the gene-expression matrix. We implemented these approaches using the NMF library in R (28).

- **Weighted non-negative Matrix Factorization (WNMF):** WNMF, also known as ls-nmf (least squares nonnegative matrix factorization) introduced by Guoli Wang et al.(22), deploys uncertainty measurements of gene expressions into NMF updating steps (22). WNMF gets a weight matrix as input to emphasize more reliable cells (m) in the factorization step. Given a non-negative matrix Rm∗n, WNMF calculates 2 nonnegative factors Um∗k and Vn∗k, minimizing a cost function as follows:

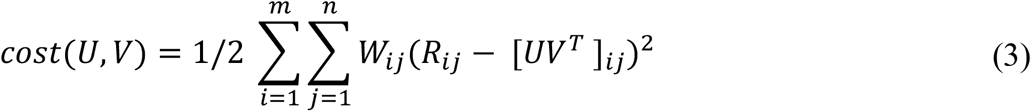
- **Sparse Non-negative Matrix Factorization (sNMF):** Sparse Non-negative Matrix Factorization introduced by Hyunsoo Kim et. al. (20), uses alternating non-negativity-constrained least squares in the updating steps, and in each step sNMF keeps the sparsity of the factorized matrices. Given a non-negative matrix Rm∗n, sNMF calculates 2 non-negative factors Um∗k and Vn∗k, which minimize the cost function as follows:

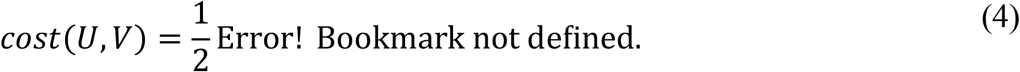

This technique is known to work well on sparse data sets, which makes it especially suitable for scRNA-seq data.

In both approaches, nndsvd is used to generate the function seed (30), and was effective in identifying a suitable initialization point as both functions converged after just 5 iterations. After factoring the gene-expression matrix with either of the above techniques, 2 non-negative factors are generated Um∗k and Vn∗k, to generate the imputed matrix:

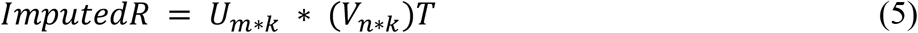

#### 2.2.3 Scaling

The second highest expression ventile (19v) encompasses multiple gene expression values near the upper end of the overall expression distribution. Such genes are less sensitive to dropout events than lower expression values and are therefore more reliable, but they also avoid extreme high-end outlier values such as erroneously high “jack-pot” values (1, 17). Thus, for gene-wise scaling to match imputed gene expression distribution to reliable data in the raw matrix, FIESTA uses the 19v, as follows:

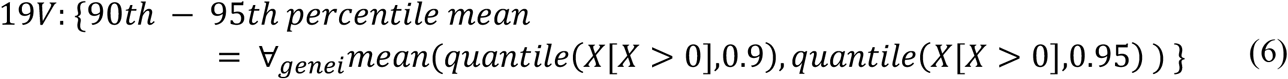

And,

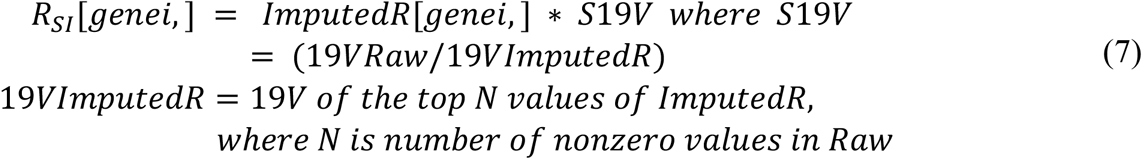

#### 2.2.4 Thresholding

The resulting imputed matrix is strictly positive, although many values are very small where expression levels are low-to-negligible in a particular cell subset or across the entire data set. Some users may wish to return extremely low imputed values to zero for purposes of visualization and interpretation. For this reason, FIESTA includes an optional post-hoc thresholding procedure for ‘zeroing-out’ predicted false positives in the imputed matrix while preserving correctly imputed values elsewhere in the data (Figure 1). First, prototypical cell state divisions are identified using the graph-based clustering approach from the Seurat package on matrix R_SI_ [seurat package clustering (31-34)].

We then identify all cluster/gene combinations that lack positive values in the input data, denoting these “null-clusters” as C_null_. We interpret these clusters to be enriched for true zeros in the raw data. By extension, their imputed values are predominantly considered to represent imputation noise/false positives. Similarly, clusters with 10 or more positive values in the raw data are identified as C_pos_, likely indicating true positives for a given gene, although within a given C_pos_ cluster the zeroes that are present may represent either true or false negatives.

Going gene-by-gene in each cluster in the imputed matrix, FIESTA estimates expression distributions by fitting a MixNormal model to each. Let X denote the expression levels of a gene within a specific cluster. We model X as a mixture of two normal distributions:

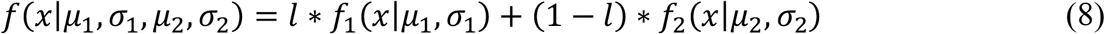

Where *f* represents the probability density function of a mix-normal distribution. μ represent the mean value and σ represents the standard deviation. *l* represents the proportion of the first normal distribution in the mixture.

This allows different subsets of cells in a cluster to have different expression distributions. Clusters that are essentially monomodal yield two overlapping fitted distributions whereas bimodal and more complex patterns will typically yield distributions with two different means and proportionality of contributing nodes. Reasoning that the majority of fit distributions in C_pos_ likely represent true positives, and conversely, the majority of lower distributions in C_null_ are true negatives, we select the midpoint between the mean of the lower distribution means for Cnull and Cpos fit distributions as a thresholding point. All distributions with means below the threshold are considered likely zero+noise distributions that can be zeroed out. Subsequently, all values in a cluster that are predicted to belong to a distribution with both means below the threshold are “zeroed-out” in the imputed matrix if they were also null in the raw data prior to imputation. On clusters where one distribution mean is above and one below the threshold, the percentage of cells predicted to belong to the lower distribution get thresholded beginning with the lowest imputed value and moving up the ranked list of values. Thus, some clusters will retain all imputed values, some will zero-out all imputed values and some will retain values corresponding to a positive distribution while zeroing values corresponding to a lower distribution depending on whether the respective distribution means are closer to C_null_ models or C_pos_ models.

## 3 Results

### 3.1 Data set-specific imputation parameters

We calculated the percentage of total variance encompassed by sequentially increasing ranks of SVD for each filtered data set, independently. For example, at k=27 in the mammary epithelial data set (17), the SVD model captured greater than 97% of the total variance and increasing k had diminishing returns (Figure 2A). We subsequently employed the 97% capture threshold for each data set we examined, resulting in unique k for each dataset analyzed. The numbers of factors recommended for each data set analyzed in the present study based on similar variance capture (i.e. 97%) is shown in Figure 2B. The lower feature identification for melanoma cell line data likely reflects the type of library preparation, less cell type heterogeneity in this data set and potentially also the cell quality as filtering cells with high mitochondrial gene content increased optimal feature resolution to k=10. The k value should ideally be tuned such that factors retain critical distinguishing features between similar cell types while optimally mitigating noise and computational burden. This parameter remains tunable by end users in the FIESTA package to allow the user to balance computation speed, and feature resolution. For consistency across data sets in these studies, we imposed an upper bound on k=25, to constrain resolution and computational load, and impose ‘low rank’ dimensionality reduction.

**Figure 2.**
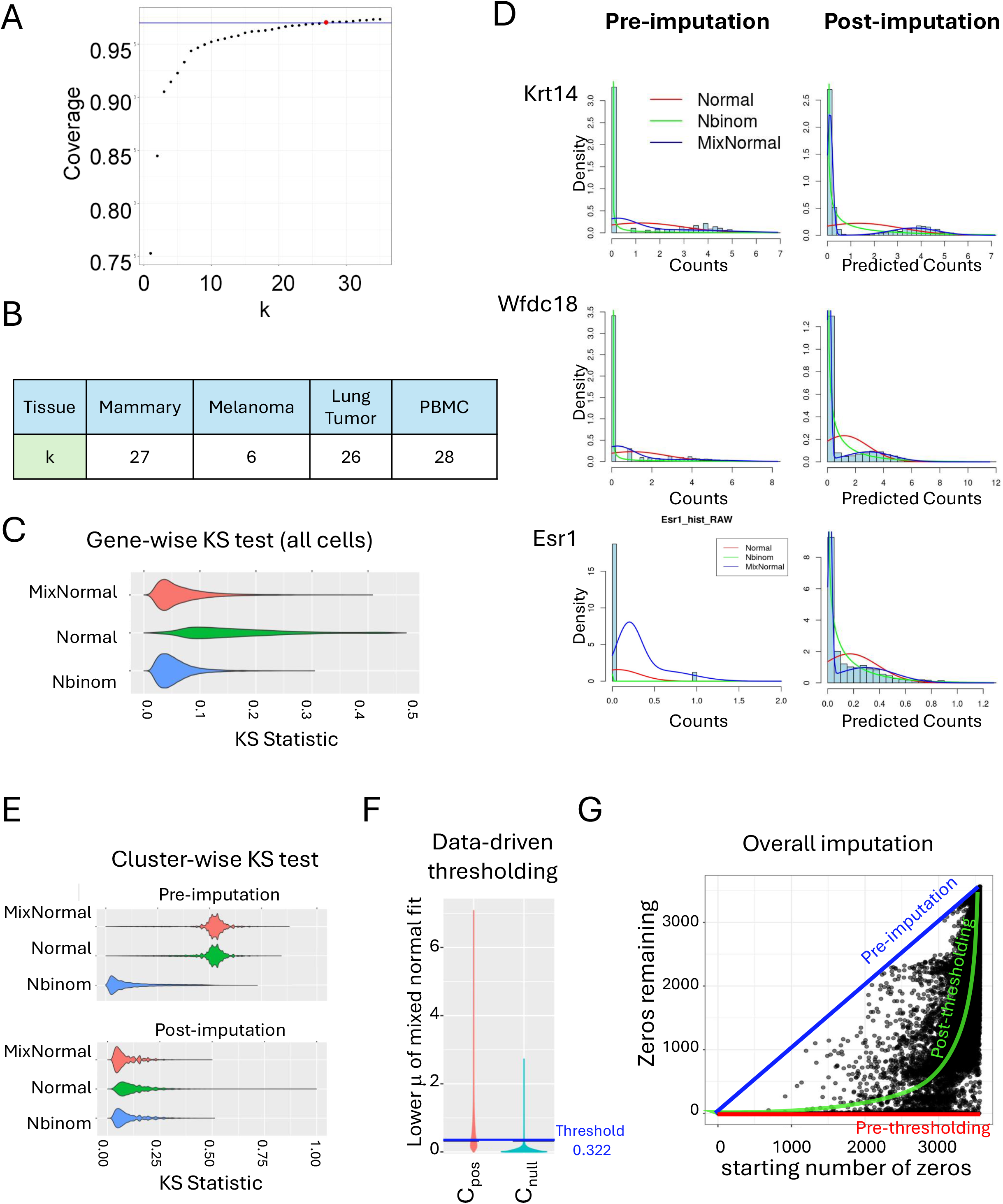
Parameter derivation from pre- and post-imputed data. A) Selection of factorization rank k from SVD diagonal matrix capture of variance in mouse mammary single cell data (17). B) Optimal k is data set dependent. C) Genewise goodness of fit to MixNormal, Normal and Negative binomial (Nbinom) distributions based on KS test statistic. D) Curve fitting on representative genes in pre-and post-imputation data. E) Genewise goodness of fit (KS statistic) of MixNormal, Normal and Nbinom distributions on data separated by cell clustering. F) Data-driven thresholding step based on lower µ from MixNormal fitting to imputed data. G) Comparison of zeroes before and after imputation and thresholding.

We applied cell-wise 19V scaling to each data set independently as described in section 2.1.3. Data sets were then factored via WNMF or sNMF and reconstituted via factor multiplication as described in section 2.2.2. Following imputation we applied gene-wise scaling using the 19V of all positive values for each gene in the raw data (19VgeneR) compared to the 19V of an equivalent number of high-end values in the imputed data (19Vgene I), to help ensure that values in the post-imputation matrices correspond to actual measured values where such values are reliable in the raw data.

Further, as described recently by Lindermann et al, many researchers wishing to recover missing values may also wish to “preserve true zeros” in the data output (3). Therefore, we devised a thresholding procedure to preserve (i.e. return to zero) the most probable true zeros within the dataset, aiming to mitigate the issue of “over-imputation”.

Poisson-gamma/negative binomial (nBinom) distribution has been commonly employed on scRNA-seq data (2). In part this may reflect the strictly non-negative but zero inflated shape of raw data. For instance after imputation, we found that a mixture of two normal models (MixNormal) and nBinom models performed similarly, whereas single normal distributions showed somewhat poorer fit to many genes (Figure 2C). Closer examination revealed that this likely reflects multimodal distributions in many genes that define distinct cell subtypes, a common feature of most modern scRNA-seq data sets (Figure 2D).

Therefore, as described in section 2.2.4, we utilized graph-based clustering approaches that are widely employed to discern cell subtypes in single cell data (31-34) and reexamined distribution fitting on clusters independently. These analyses indicate that MixNormal distributions provide reasonable models of imputed gene expression for most genes following imputation and clustering (Figure 2E). This likely reflects the ability of MixNormal distributions to fit both bimodal and monomodal distributions by returning two largely overlapping normal distributions in the latter case (Figure 2F and Supplemental Figure 1). Furthermore, in cases where the fit of the normal distribution with the lower mean is in the range of C_null_ the approach is consistent with the standard practice of modeling noise with a normal distribution.

As expected, C_null_ distributions were distinguishable from C_pos_ distributions allowing for automatic threshold identification (Figure 2F). Although application of post hoc thresholding returned many imputed values to zero, the majority of these had extremely low positive values in the imputed data already. Furthermore, following the application of these scaling and thresholding steps, which restore many zeros in the data, the imputation pipeline still noticeably reduced the number of zero values present in the expression matrix relative to the non-imputed (Figure 2G). Hence, this supports the interpretation that their true expression (both prior to and after imputation) is negligible. However, it also raises questions about the overall value of thresholding on data interpretation, a topic which has been raised previously (see published reviews accompanying ref. 3) and which we evaluate in greater depth below.

### 3.2 FIESTA’s imputation accuracy compares favorably with alternative approaches across a broad range of expression values

Comparing the overall level of imputation and preservation of zeros between FIESTA and several other imputation approaches including ALRA, MAGIC and SAVER, we find that FIESTA exhibits a higher imputation rate than ALRA while preserving more zeros than either SAVER or MAGIC (Figure 3A-C). For instance, we examined predicted expressions in human PBMC data sets that were similarly analyzed by Linderman et al. (3), and which have field knowledge that can inform interpretation of imputation/thresholding accuracy. Respectively, 78%, 74% and 74% of expected zeros were returned as zeros following thresholding of FIESTA-imputed PBMC data for B-cells, T-cells and Monocytes (Figure 3A). Conversely, thresholded FIESTA-imputed data retained 71%, 74 % and 70% of imputed positive values that had been zero in the input data for these cells. Using the nomenclature of Lindermann et al., retained zeros can be referred to as ‘zeros preserved’ (ZP) and retained positive imputed values that were zero in the input data can be referred to as total zero completed (TC) (3). In comparison with the FIESTA derived values above, ALRA was conservative (ZP: B-cells 93%, T-cells: 92%, Monocytes: 94%, TC: B-cells 57%, T-cells 56%, Monocytes 53%); Magic was liberal (ZP: B-cells 59%, T-cells 37%, Monocytes 38%, TC: B-cells 85%, T-cells 92%, Monocytes 88%); and Saver was more balanced (ZP: B-cells 78%, T-cells 77%, Monocytes 73%, TC: B-cells 69%, T-cells 69%, Monocytes 70%), though Saver was slightly more conservative than thresholded FIESTA (Figure 3A). However, it should also be mentioned that the ‘ground truth’ for ZP and TC metrics relies on discriminatory gene expression that could be dogmatic as complete absence of expression in a presumed “negative” cell population is often unconfirmed. Indeed, when looking at expected discriminatory genes between these cell types in HUMAN cell atlas data (35) many scenarios are observed where ‘negative’ genes for a particular cell type, are, in reality, expressed at a low level (e.g. NCAM1, CD8a, CD4, CD70) (Supplemental Figure 2). Thus, we also found it critical to assess whether values are *properly* thresholded by cell type, since higher ZP estimates could be a trivial result of reduced overall imputation.

**Figure 3.**
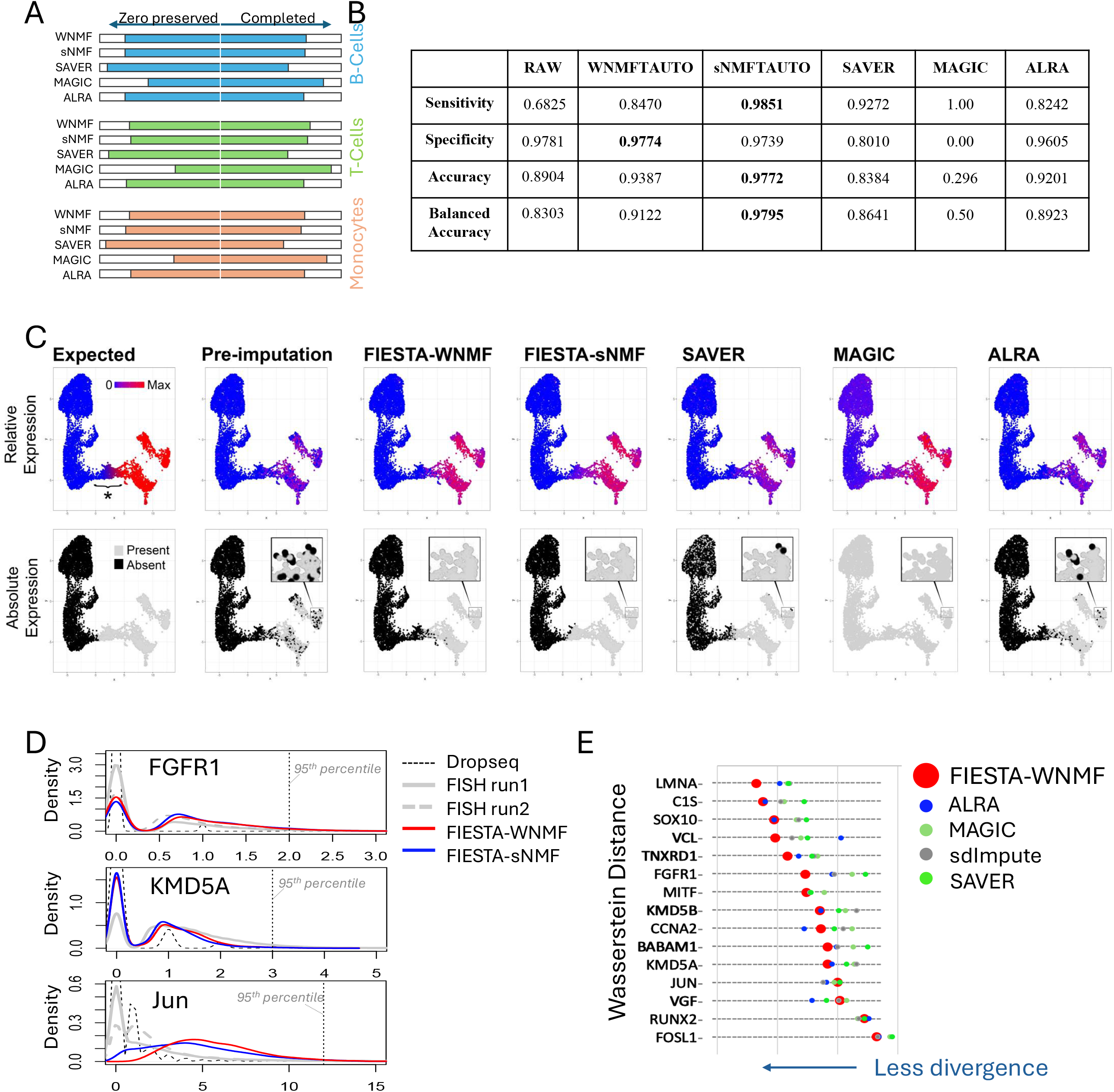
Performance of FIESTA in data recovery compared to existing methods. A) Ratio of biological zeros preserved (ZP) alongside to total zeros completed (TC) over three distinct cell types, B-cells, T-cells and Monocytes (23). B) Sensitivity, specificity, and Accuracy measures for imputed Nkx2.1 in Zewdu et al. (24) C) Graphical representation (umap) of imputation accuracy and identification of Nkx2.1 positive cells in Zewdu et al. by various imputation approaches with ground truth based on genetic deletion of the Nkx2.1 gene. D) Representative distributions of gene expression by Dropseq (i.e. pre-imputation), FISH (orthogonal analysis technique) and FIESTA imputation on melanoma cells (2,23). E) Wasserstein distance metric comparing FIESTA to other approaches and to FISH in melanoma cells (2,23).

As an additional test of FIESTA’s accuracy of imputation and thresholding in this regard, we leveraged a single cell data set derived from a mouse lung adenocarcinoma study where knowledge of the genetic system provides a surrogate ground truth (24). This work involved experimentally induced genetic recombination to delete the Nkx2.1 gene. We inferred that although Nkx2.1 transcripts were detected in only a subset of cells where gene deletion was not experimentally activated, it was likely expressed but undetected as it is critical in the phenotype of co-clustered cells that share the Nkx2.1 positive phenotype (24). Using this inferred Nkx2.1 expression pattern, we tested the sensitivity, specificity, and balanced accuracy of FIESTA vs. various alternative imputation approaches (Figure 3B). In this context, MAGIC was overly liberal resulting in a strictly positive expression for Nkx2.1, thus exhibiting 100% sensitivity but no calculable specificity in this analysis (Figure 3B, C). The remaining approaches were more balanced with Saver being somewhat more liberal and ALRA being more conservative (Figure 3B, C). The overall highest sensitivity, specificity and accuracy was achieved by FIESTA using sNMF (Figure 3B, C).

Additionally, a number of prior studies have leveraged orthogonal data from single molecule fluorescence in situ hybridization (smFISH) in a melanoma cell line data set -- an alternate approach to quantification of transcript abundance in individual cells, -- to serve as a reasonable estimation of the true relative expression, and therefore a surrogate ground truth (2, 3). With this data set we may compare values recommended by FIESTA with abundance of 15 distinct, variable transcripts in individual melanoma cells by smFISH that were also assessed by parallel scRNA-seq from the same cell populations (23). We observed that FIESTA effectively restored relative expression levels, aligning more closely with FISH data in comparison with raw values acquired from Dropseq (Figure 3D, E). In most cases, FIESTA had better performance over alternative approaches (Figure 3E). Nevertheless, we urge caution in using these data sets as we identified strong batch effects for some of the reported genes (Supplemental Figure 3) and were unable to find matched control data demonstrating specificity of the probes used.

### 3.3 Imputation affects gene-cell and gene-gene relationships

We applied FIESTA to adult mouse mammary gland scRNA-seq profiles that we published previously (17). In this mammary gland data set, we previously described three major cell types in detail based on transcriptional profiles and clustering (17). These cell types correspond to long established differentiated cell types of the mammary gland with distinguishing marker gene expression patterns, including Krt14 for the basal cell type, Wfdc18 for the alveolar cell type and Krt18 (in the absence of Wfdc18) for the remainder of luminal cells including the hormone sensing epithelial cells of the mammary gland (17, 36). Annotating cells of each major cell type, examination of several factors known to mark distinct cell types in the mammary gland revealed expression patterns in scRNA-seq data that were too sparse and digitized for straightforward visual interpretation (Figure 4A). FIESTA imputation restored anticipated cell type specific expression patterns for each of the factors we examined (Figure 4A-C). For instance, FIESTA identified Epcam^High^Itga6^Low^ and Epcam^Low^Itga6^High^ cell types consistent with well-known expression patterns of their gene products in different compartments of the mouse mammary epithelium, and similarly enabled discernment of Ly6A (sca1) high and low populations within the epithelium consistent with distinct cell states described in prior reports (Figure 4B) (37-40). Conservative imputation approaches left these genes and their cognate relationships highly digitized, while other approaches were overly liberal or showed arbitrary thresholding effects that do not well match the relationships described for their protein products in the literature (Supplemental Figure 4). Imputation was cell type specific, predicting expression of estrogen receptor (Esr1) strictly in non-alveolar luminal cells and expression of the basal transcription factor TP63 strictly in basal cells (Figure 4C) (41).

**Figure 4.**
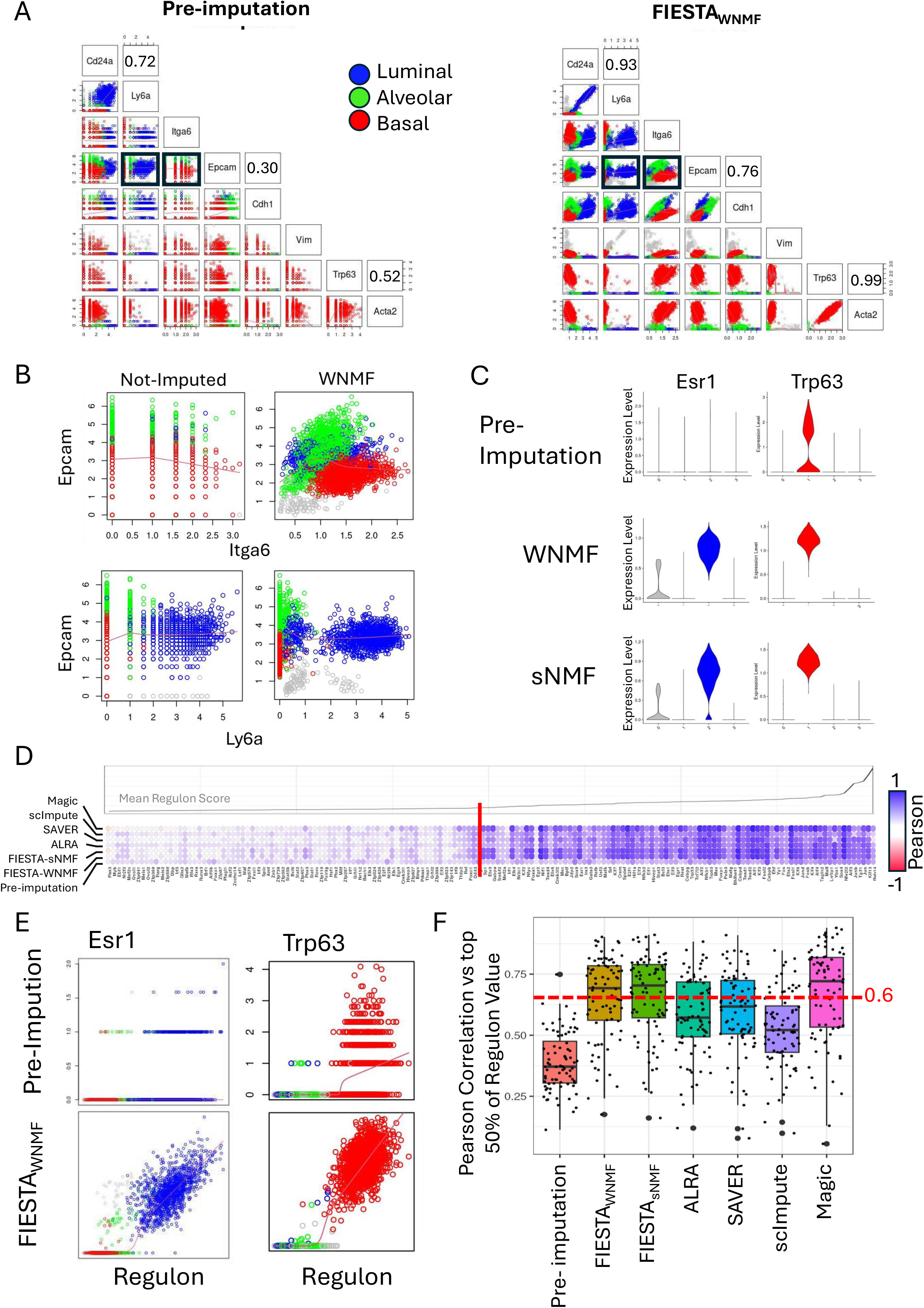
Correlation of FIESTA imputation values with orthogonal gene expression predictions. A) Mouse mammary cell-type marker-gene correlations before and after imputation. B) Pre- and post-imputation co-expression plot for Epcam and Ly6a or Itga6.C) Cell type specificity of FIESTA imputed Esr1 and Trp63 values compared to pre-imputation levels. D) Correlation of imputed transcription factor values with Regulon activity scores given by SCENIC (41) in the Giraddi et al. (17). The top half of regulon scores show high correlation with imputed values from several approaches including FIESTA. E) Plot of pre-imputation (upper panels) and FIESTA imputed values (lower panels) for Esr1 and Trp63 vs SCENIC regulon scores F) FIESTA has the highest proportion of strong and very strong (i.e. Pearson>0.6) correlations with regulon predictions from SCENIC among imputation approaches.

Transcription factors (TF) are key regulators of cell state but are notoriously problematic in scRNA-seq data as they are often lowly expressed and prone to dropout. To gauge TF imputation accuracy, we compared FIESTA imputed values to regulon values calculated by SCENIC (17, 42). SCENIC uses correlated genes across a data set as well as transcription factor binding site information to identify whether a given transcription factor is likely to be active. Where SCENIC predicts transcription factor activity it can be inferred that the transcription factor was present even if it was not well-detected in the raw scRNA-seq data. Thus, although only applicable to transcription factors, SCENIC essentially provides an orthogonal imputation approach for active transcription factors. FIESTA demonstrated good correlation of imputed transcription factor values with regulons in the upper half (i.e. active) subset of regulons (Figure 4D-F). For instance, the luminal transcription factor Esr1 and the basal transcription factor Trp63 showed very strong positive correlations between the SCENIC predicted activity level and the FIESTA predicted expression level despite being highly digitized in the raw data (Figure 4E). For comparison, we also imputed the same data set using other published approaches and assessed their correlations to SCENIC regulon values as well (Figure 4D, F). FIESTA showed the best overall performance with over 73.7% of FIESTA imputed transcription factors for active regulons showing strong to very strong correlation (r^2^>0.6). MAGIC also showed high correlation with 68.7% of TF showing strong correlation followed by SAVER (48.7%), ALRA (47.5%), and scImpute (7.5%). As, the lower half of predicted SCENIC scores can equally represent absence of the underlying transcription factor *or* its inactivity through post transcriptional control, they are not used in this comparison. Enhanced detection of lowly expressed transcription factors by FIESTA was also observed in other data sets (24).

### 3.4 Imputation and thresholding enable identification of additional differentially expressed genes

Differential expression analysis on non-imputed data using Wilcoxon Rank Sum test in Seurat (34, 43-45) identified 295, 691, and 344 differentially expressed genes (DEG) uniquely overexpressed in basal vs Alveolar, Basal vs luminal and Luminal vs alveolar cellular subtypes, respectively (Figure 5A). These genes represent many well-known and often robustly expressed cell type identifiers in the mammary epithelium. We next asked how FIESTA imputation would affect identification of DEGs. We initially found that FIESTA greatly expanded the list of DEG detected by Seurat (Figure 5A and Supplemental Figure 5B). Within the expanded gene list provided by imputation were genes with clearly cell-type restricted gene expression patterns but whose expression was detected only in a restricted number of cells in the raw data (Figure 5B). Thus, FIESTA helps identify DEG candidates for downstream analysis that would have been missed in traditional scRNA-Seq analysis including sparsely represented, lowly expressed cell type markers and transcription factors exhibiting lower expression levels with comparatively large biological effects (Supplementary Figure 5A,B). Retention of most DEG from raw data, extension of the DEG candidate list by imputation, concurrence between FIESTA (using either sNMF or WNMF) and ALRA were all confirmed using PBMC data (Supplementary Figure 5C).

**Figure 5.**
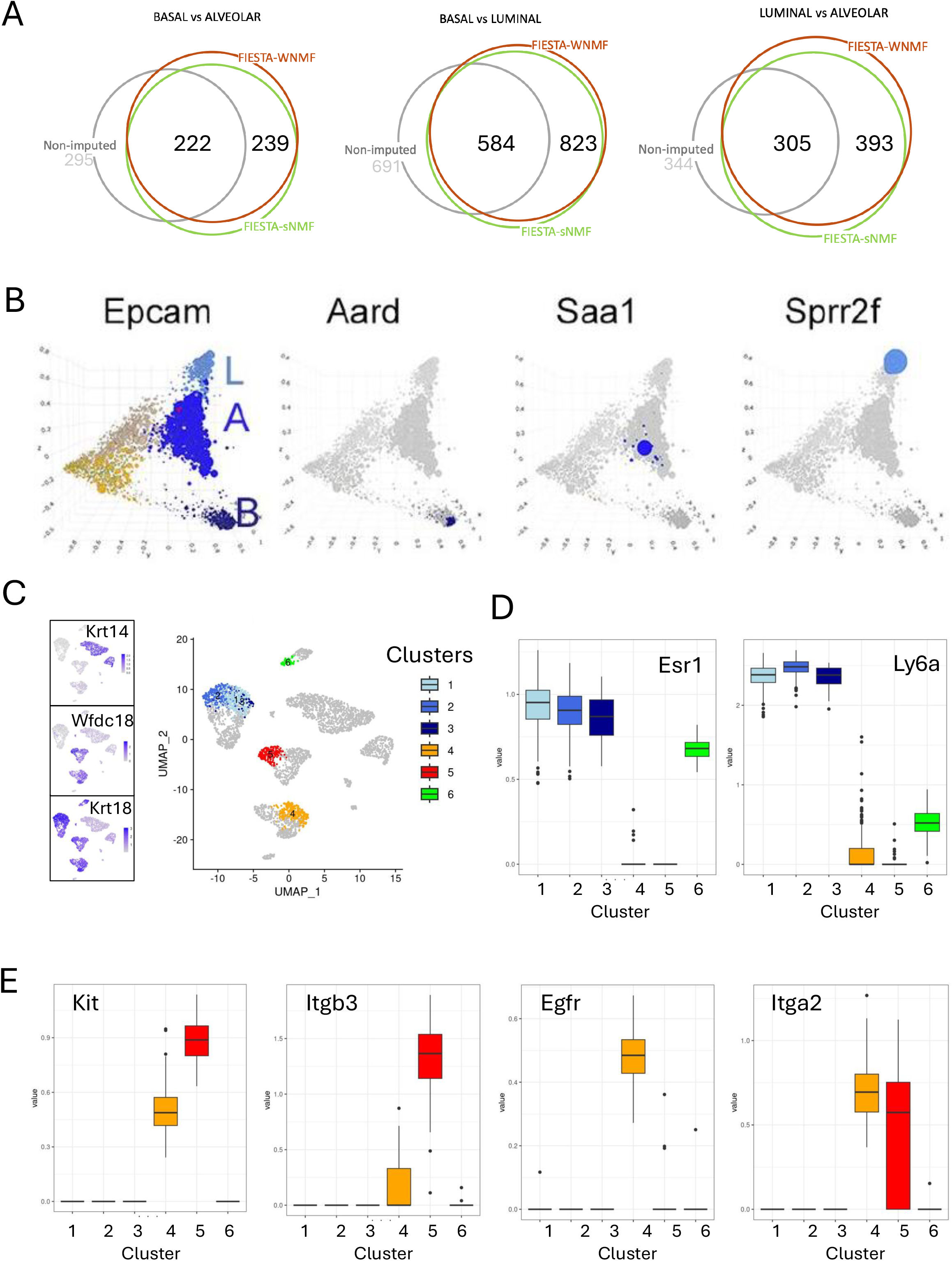
Identification of novel cell type markers from imputed data. A) Differentially expressed genes identified in pre-imputation data vs FIESTA imputed data. Diffusion map (as in Giraddi et al.) showing cell type specificity of genes specifically detected as differentially expressed following FIESTA imputation. C-E) Mouse mammary cellular subtypes implied by imputed data. C) UMAP and select cell sub-clusters. D-Identification of Esr1^high^Ly6a^high^ and Esr1^high^Ly6a^Low^ subtypes among luminal cells. E) Identification of Kit^high^Itgb3^high^ and Egfr^high^Itga2^high^ alveolar cell suptypes.

### 3.5 Imputation affects cell-cell relationships

While all approaches retained high classification accuracy for the major cell types identified in the adult mouse mammary data set, we examined whether discrete (sub-)cell types could be identified following imputation. Examining two presumptive alveolar and two presumptive luminal cell subsets based on UMAP coordinates and expression of transcripts related to known surface markers in the mammary gland, we identified distinguishable Egfr+Itga2+ and Kit+Itgb3+ positive subsets of Wfdc18+ cells, and also identified a Ly6a-Low subset of Esr1+ cells (Figure 5C, D). Although these cellular sub-states were not readily distinguishable in the non-imputed data, they may correspond to select cellular sub-states emerging from other recently published studies of cellular sub-types in the mammary gland (46-48).

Differential gene expression analysis between these subsets revealed that the cells are likely distinguishable by broader transcriptomic differences representing distinct (although still quite similar) cell states (Figure 5D).

## 4 Discussion

In this work, we set out to examine the potential of matrix factorization via NMF and subsequent matrix completion from factor multiplication to accurately impute missing values in scRNA-seq data. This effort was based on the demonstrated effectiveness of NMF in identifying meaningful cell and gene relationships in gene expression data sets, and its proven utility in recommender systems dealing with other types of sparse data matrices. By applying unsupervised feature selection, scaling and thresholding parameters estimated from measured values in the raw data in conjunction with imputed value predictions, we produced an unsupervised computational imputation pipeline for data processing, FIESTA, that outperforms several alternative approaches. Further, by comparing results of this approach to expression patterns known from the literature and orthogonal analysis in four distinct published data sets, we found FIESTA produced reliable relative expression patterns for many genes including those poorly detected in the raw data.

Recently, Lin and Boutros also demonstrated the inherent potential for NMF based matrix completion to impute missing values in expression data sets and suggested several approaches for algorithm improvement that may reduce computational burden and increase speed (49). Indeed, computational burden has been the major challenge to scalability of NMF (29). However, recent advances promise significant improvements for scalability (29, 50). Whether through increased speed or iterative application to sequential subsets of the data, NMF approaches such as FIESTA will likely have useful application to many of the emerging data sets and compendia that have been produced with the scRNA-seq technology (e.g. (35, 51)). Further, as imputation need only be carried out once per data set, and once complete incurs no further computational burden on downstream analysis, the incorporation of NMF in the FIESTA pipeline provides an acceptable tradeoff for imputation accuracy even prior to improvements in scalability. In contrast, despite their speed and fairly reasonable gene expression estimates for many genes, some existing methods may be overly conservative (e.g. Saver and ALRA), too liberal (e.g. MAGIC), or require potentially biased priors in the form of assumptions about ‘important’ genes or expected cell types. In contrast, FIESTA is unsupervised and gives expression values that realistically reflect both cellular heterogeneity and lineage/cell-type restriction.

We also recognize that ‘gene pulsing’ may also contribute to null entries observed in raw data (52, 53) and in this regard imputed data such as that produced by FIESTA may be viewed as integral or mean expression models for such genes, relating its generalized expression to the cell type and larger gene circuit in which they operate. Additionally, although the large number of differentially expressed genes modeled by imputation of sparse scRNA-seq data may contain some false positives (54), it also permits the identification of novel candidate cellular subtypes and markers for future study that were missed by the same analysis of non-imputed data.

In summary, FIESTA, which is based on feature identification using WNMF or sNMF, imputes missing values in scRNA-seq data with a more intuitive relationship to known biology than previously reported approaches. While all approaches tested altered graphical representation of the data and thus affect interpretability of the data including cell-cell and cell-gene relationships, the expression patterns following FIESTA worked across a broad range of input values and detection frequencies, and matched values obtained from orthogonal approaches. Analysis of gene expression differences among populations emerging from FIESTA suggests resolution of biologically meaningful features and an enhanced ability to detect differentially expressed genes that define minority cell types. The current version of FIESTA (v 1.2) as described in this manuscript is freely and immediately available as an R package [https://github.com/TheSpikeLab/FIESTA].

## 5 Supporting information

Supplemental Figure 1: Examples of mixed normal fitting on a Krt14 and an Esr1 expressing cluster and on a sampled normal distribution.

Supplemental Figure 2: Graphical representation of single cell expression data from the Human Protein Atlas (34), showing sporadic expression of transcripts in presumed negative populations.

Supplemental Figure 3: Examples of three genes in FISH data (23) that show strong batch effects.

Supplemental Figure 4: Predicted (i.e. imputed) expression levels of known mouse mammary cell type markers following various imputation approaches.

Supplemental Figure 5: Comparison of differentially expressed gene levels before and after FIESTA imputation.

## References

1. Marinov GK, Williams BA, McCue K, Schroth GP, Gertz J, Myers RM, Wold BJ. From single-cell to cell-pool transcriptomes: Stochasticity in gene expression and RNA splicing. Genome Research. 2014;24(3):496–510.

2. Huang M, Wang J, Torre E, Dueck H, Shaffer S, Bonasio R, et al. SAVER: gene expression recovery for single-cell RNA sequencing. Nature Methods. 2018;15(7):539–42.

3. Linderman GC, Zhao J, Roulis M, Bielecki P, Flavell RA, Nadler B, Kluger Y. Zero-preserving imputation of single-cell RNA-seq data. Nature Communications. 2022;13(1).

4. Li WV, Li JJ. An accurate and robust imputation method scImpute for single-cell RNA-seq data. Nature Communications. 2018;9(1).

5. Arisdakessian C, Poirion O, Yunits B, Zhu X, Garmire LX. DeepImpute: an accurate, fast, and scalable deep neural network method to impute single-cell RNA-seq data. Genome Biology. 2019;20(1).

6. Van Dijk D, Sharma R, Nainys J, Yim K, Kathail P, Carr AJ, et al. Recovering Gene Interactions from Single-Cell Data Using Data Diffusion. Cell. 2018;174(3):716–29.e27.

7. Xu J, Cai L, Liao B, Zhu W, Yang J. CMF-Impute: an accurate imputation tool for single-cell RNA-seq data. Bioinformatics. 2020;36(10):3139–47.

8. Ronen J, Akalin A. netSmooth: Network-smoothing based imputation for single cell RNA-seq. F1000Research. 2018;7.

9. Zand M, Ruan J. Network-based single-cell rna-seq data imputation enhances cell type identification. Genes. 2020;11(4):377.

10. Chen M, Zhou X. VIPER: variability-preserving imputation for accurate gene expression recovery in single-cell RNA sequencing studies. Genome biology. 2018;19(1):196.

11. Tang W, Bertaux F, Thomas P, Stefanelli C, Saint M, Marguerat S, Shahrezaei V. bayNorm: Bayesian gene expression recovery, imputation and normalization for single-cell RNA-sequencing data. Bioinformatics. 2020;36(4):1174–81.

12. Ling Q, Xu Y, Yin W, Wen Z, editors. Decentralized low-rank matrix completion 2012: IEEE.

13. Cai J-F, Candès EJ, Shen Z. A Singular Value Thresholding Algorithm for Matrix Completion. SIAM Journal on Optimization. 2010;20(4):1956–82.

14. Dai X, Zhang N, Zhang K, Xiong J. Weighted Nonnegative Matrix Factorization for Image Inpainting and Clustering. International Journal of Computational Intelligence Systems. 2020;13(1):734.

15. Brunet J-P, Tamayo P, Golub TR, Mesirov JP. Metagenes and molecular pattern discovery using matrix factorization. Proceedings of the national academy of sciences. 2004;101(12):4164–9.

16. Zhu X, Ching T, Pan X, Weissman SM, Garmire L. Detecting heterogeneity in single-cell RNA-Seq data by non-negative matrix factorization. PeerJ. 2017;5:e2888.

17. Giraddi R, Chung C, Heinz R, Balcioglu O, Novotny M, Trejo C, et al. Single-cell transcriptomes distinguish stem cell state changes and lineage specification programs in early mammary gland development. Cell Rep 24 (6): 1653–1666. e1657. 2018.

18. Marjanovic ND, Hofree M, Chan JE, Canner D, Wu K, Trakala M, et al. Emergence of a high-plasticity cell state during lung cancer evolution. Cancer cell. 2020;38(2):229–46. e13.

19. Gillis N, Glineur F. Low-Rank Matrix Approximation with Weights or Missing Data Is NP-Hard. SIAM Journal on Matrix Analysis and Applications. 2011;32(4):1149–65.

20. Kim H, Park H. Sparse non-negative matrix factorizations via alternating non-negativity-constrained least squares for microarray data analysis. Bioinformatics. 2007;23(12):1495–502.

21. Kim Y-D, Choi S, editors. Weighted nonnegative matrix factorization. 2009 IEEE international conference on acoustics, speech and signal processing; 2009: IEEE.

22. Wang G, Kossenkov AV, Ochs MF. LS-NMF: A modified non-negative matrix factorization algorithm utilizing uncertainty estimates. BMC Bioinformatics. 2006;7(1):175.

23. Torre E, Dueck H, Shaffer S, Gospocic J, Gupte R, Bonasio R, et al. Rare Cell Detection by Single-Cell RNA Sequencing as Guided by Single-Molecule RNA FISH. Cell Systems. 2018;6(2):171–9.e5.

24. Zewdu R, Mehrabad EM, Ingram K, Fang P, Gillis KL, Camolotto SA, et al. An NKX2-1/ERK/WNT feedback loop modulates gastric identity and response to targeted therapy in lung adenocarcinoma. Elife. 2021;10:e66788.

25. Zheng GX, Terry JM, Belgrader P, Ryvkin P, Bent ZW, Wilson R, et al. Massively parallel digital transcriptional profiling of single cells. Nature communications. 2017;8(1):14049.

26. Peter R, Shivapratap G, Divya G, Soman K. Evaluation of SVD and NMF methods for latent semantic analysis. International Journal of Recent Trends in Engineering. 2009;1(3):308.

27. Phillips RD, Watson LT, Wynne RH, Blinn CE. Feature reduction using a singular value decomposition for the iterative guided spectral class rejection hybrid classifier. ISPRS Journal of Photogrammetry and Remote Sensing. 2009;64(1):107–16.

28. Gaujoux R, Seoighe C. A flexible R package for nonnegative matrix factorization. BMC bioinformatics. 2010;11:1–9.

29. Moon GE, Ellis JA, Sukumaran-Rajam A, Parthasarathy S, Sadayappan P, editors. ALO-NMF: Accelerated locality-optimized non-negative matrix factorization. Proceedings of the 26th ACM SIGKDD International Conference on Knowledge Discovery & Data Mining; 2020.

30. Boutsidis C, Gallopoulos E. SVD based initialization: A head start for nonnegative matrix factorization. Pattern recognition. 2008;41(4):1350–62.

31. Xu C, Su Z. Identification of cell types from single-cell transcriptomes using a novel clustering method. Bioinformatics. 2015;31(12):1974–80.

32. Levine JH, Simonds EF, Bendall SC, Davis KL, El-ad DA, Tadmor MD, et al. Data-driven phenotypic dissection of AML reveals progenitor-like cells that correlate with prognosis. Cell. 2015;162(1):184–97.

33. Blondel VD, Guillaume J-L, Lambiotte R, Lefebvre E. Fast unfolding of communities in large networks. Journal of statistical mechanics: theory and experiment. 2008;2008(10):P10008.

34. Hao Y, Stuart T, Kowalski MH, Choudhary S, Hoffman P, Hartman A, et al. Dictionary learning for integrative, multimodal and scalable single-cell analysis. Nature biotechnology. 2024;42(2):293–304.

35. Regev A, Teichmann SA, Lander ES, Amit I, Benoist C, Birney E, et al. The human cell atlas. elife. 2017;6:e27041.

36. Bach K, Pensa S, Grzelak M, Hadfield J, Adams DJ, Marioni JC, Khaled WT. Differentiation dynamics of mammary epithelial cells revealed by single-cell RNA sequencing. Nature communications. 2017;8(1):1–11.

37. Shehata M, Teschendorff A, Sharp G, Novcic N, Russell IA, Avril S, et al. Phenotypic and functional characterisation of the luminal cell hierarchy of the mammary gland. Breast cancer research. 2012;14:1–19.

38. Spike BT, Engle DD, Lin JC, Cheung SK, La J, Wahl GM. A mammary stem cell population identified and characterized in late embryogenesis reveals similarities to human breast cancer. Cell stem cell. 2012;10(2):183–97.

39. Asselin-Labat M-L, Vaillant F, Shackleton M, Bouras T, Lindeman G, Visvader J, editors. Delineating the epithelial hierarchy in the mouse mammary gland. Cold Spring Harbor symposia on quantitative biology; 2008: Cold Spring Harbor Laboratory Press.

40. Balcioglu O, Gates BL, Freeman DW, Hagos BM, Mehrabad EM, Ayala-Talavera D, Spike BT. Mcam stabilizes a luminal progenitor-like breast cancer cell state via Ck2 control and Src/Akt/Stat3 attenuation. bioRxiv. 2023:2023.05.10.540211.

41. Zeps N, Bentel JM, Papadimitriou JM, D’Antuono MF, Dawkins HJ. Estrogen receptor-negative epithelial cells in mouse mammary gland development and growth. Differentiation. 1998;62(5):221–6.

42. Aibar S, González-Blas CB, Moerman T, Huynh-Thu VA, Imrichova H, Hulselmans G, et al. SCENIC: single-cell regulatory network inference and clustering. Nature methods. 2017;14(11):1083–6.

43. Korsunsky I, Nathan A, Millard N, Raychaudhuri S. Presto scales Wilcoxon and auROC analyses to millions of observations. BioRxiv. 2019:653253.

44. Hao Y, Hao S, Andersen-Nissen E, Mauck WM, Zheng S, Butler A, et al. Integrated analysis of multimodal single-cell data. Cell. 2021;184(13):3573–87. e29.

45. Stuart T, Butler A, Hoffman P, Hafemeister C, Papalexi E, Mauck WM, et al. Comprehensive integration of single-cell data. cell. 2019;177(7):1888–902. e21.

46. Regan JL, Smalley MJ. Integrating single-cell RNA-sequencing and functional assays to decipher mammary cell states and lineage hierarchies. NPJ Breast Cancer. 2020;6(1):32.

47. Fu NY, Nolan E, Lindeman GJ, Visvader JE. Stem cells and the differentiation hierarchy in mammary gland development. Physiological reviews. 2020.

48. Pervolarakis N, Nguyen QH, Williams J, Gong Y, Gutierrez G, Sun P, et al. Integrated single-cell transcriptomics and chromatin accessibility analysis reveals regulators of mammary epithelial cell identity. Cell reports. 2020;33(3).

49. Lin X, Boutros PC. Optimization and expansion of non-negative matrix factorization. BMC bioinformatics. 2020;21(1):7.

50. DeBruine ZJ, Melcher K, Triche Jr TJ. Fast and robust non-negative matrix factorization for single-cell experiments. BioRxiv. 2021:2021.09.01.458620.

51. Consortium TM, twc@ stanford. edu 4 5 6 g Darmanis Spyros spyros. darmanis@ czbiohub. org 2 h OcSNKJNNFMAPQSRqsefW-CT, Oliveira 2 Sit Rene V. 2 Stanley Geoffrey M. 3 Webber James T. 2 Zanini Fabio 3 LcBJBOCMBCSGFJRCMAPLPA, Stanley CdaBJBOCPCDDSDJLKJPAO. Single-cell transcriptomics of 20 mouse organs creates a Tabula Muris. Nature. 2018;562(7727):367–72.

52. Chubb JR, Trcek T, Shenoy SM, Singer RH. Transcriptional pulsing of a developmental gene. Current biology. 2006;16(10):1018–25.

53. Eck E, Moretti B, Schlomann BH, Bragantini J, Lange M, Zhao X, et al. Single-cell transcriptional dynamics in a living vertebrate. bioRxiv. 2024.

54. Andrews TS, Hemberg M. False signals induced by single-cell imputation. F1000Research. 2018;7.

