## Supplemental Figures 1-5 for "Factorization-based Imputation of Expression in Single-cell Transcriptomic Analysis (FIESTA) recovers Gene-Cell-State relationships"

Supplemental FIGURE 1

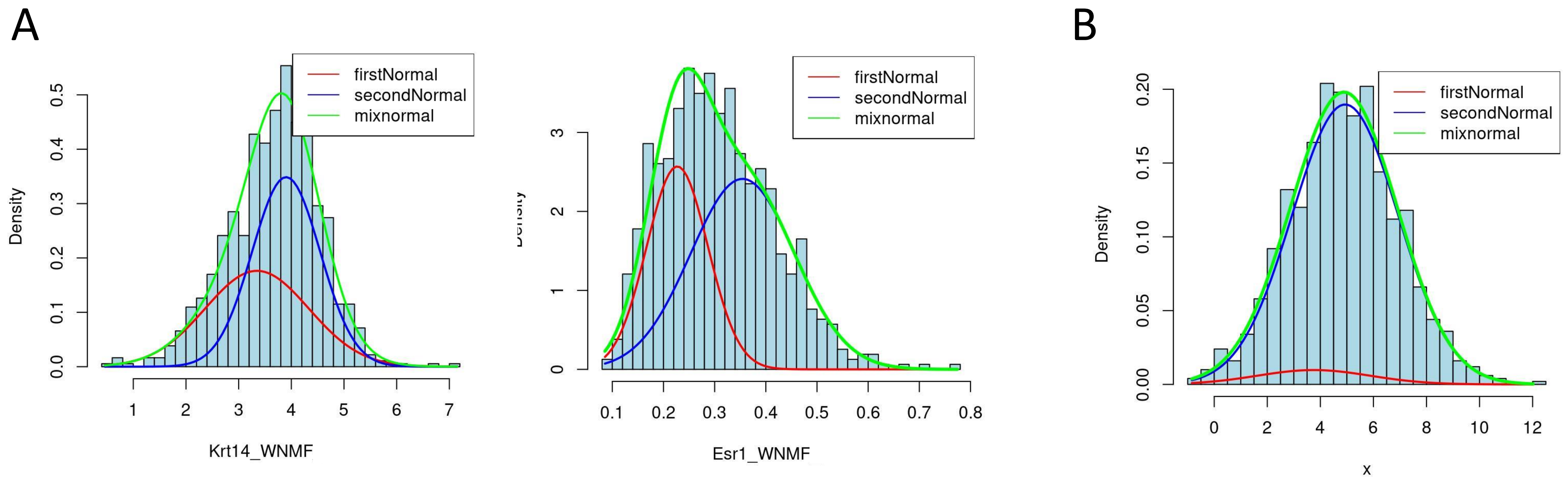


**Supplemental Figure 1.** A) Mixed normal distribution fitting for individual clusters of cells expressing keratin 14 (left) and estrogen receptor (right) following FIESTA imputation. B) A sampled normal distribution fit with a mixed normal. The area under each curve is dictated by the $l$ estimation of its proportional contribution to the mixed model.

Supplemental FIGURE 2

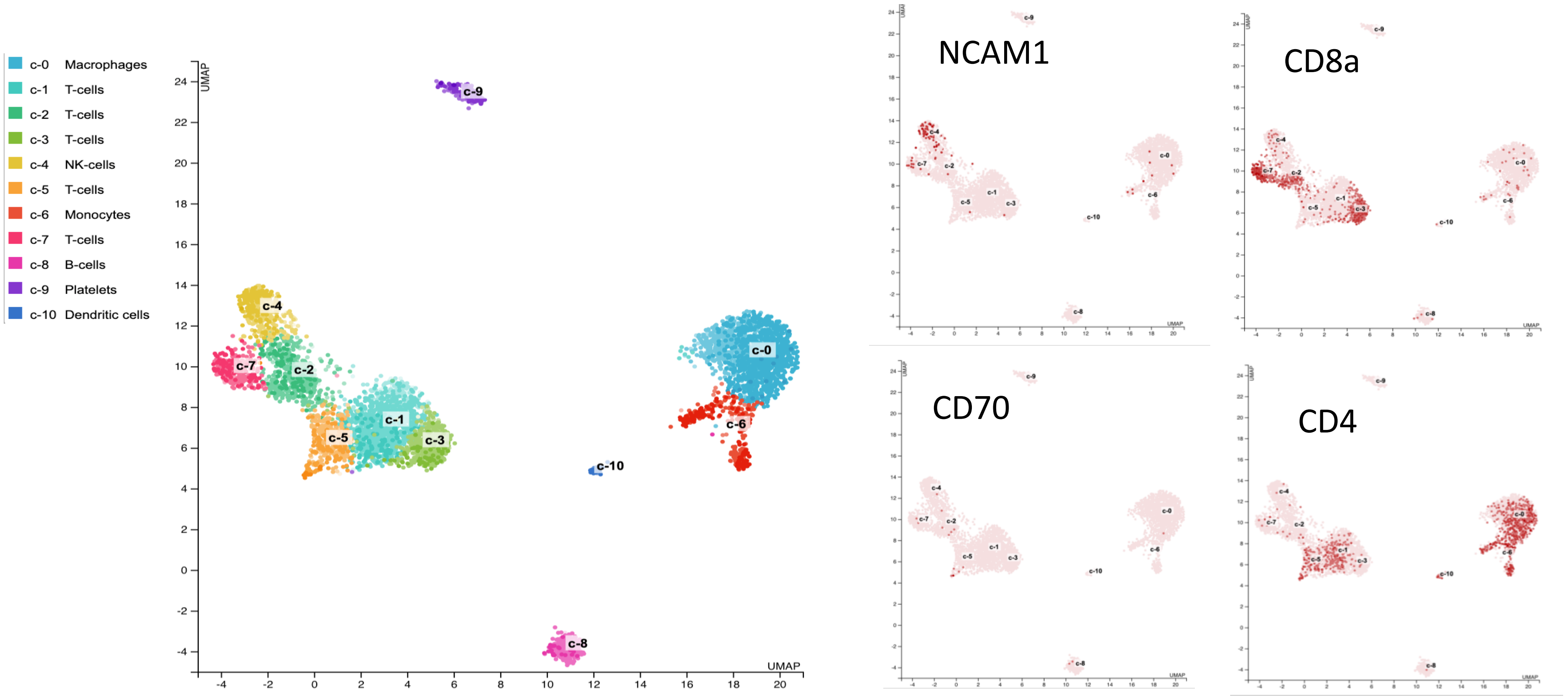


**Supplemental Figure 2.** UMAP of PBMC data (25) with clusters denoting specific cell types (left), and expression of presumed cell type markers (right).

Supplemental FIGURE 3

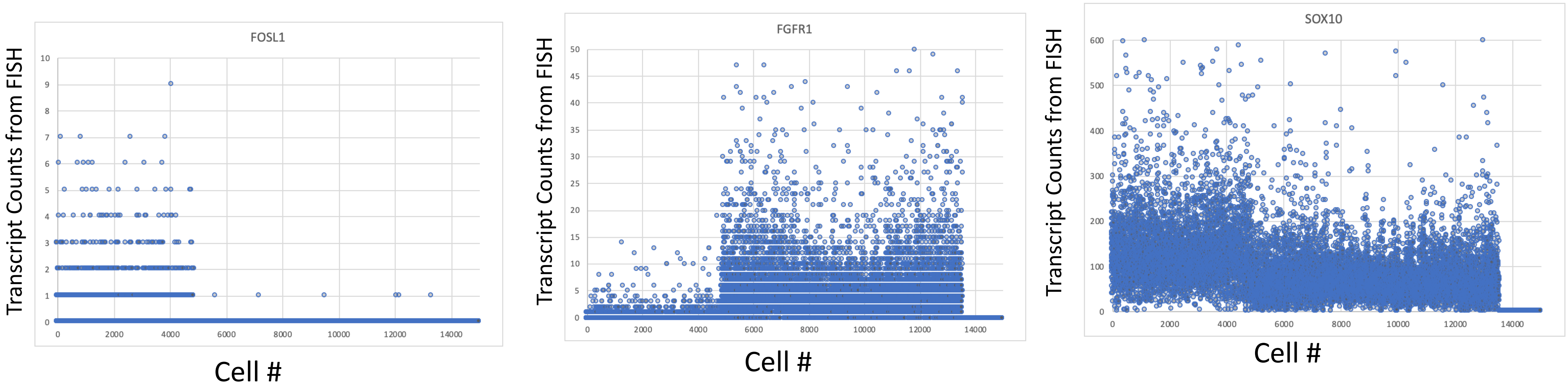


**Supplemental Figure 3.** Transcript counts in Melanoma FISH data (23) for three genes (FOSL1, FGFR1, and SOX10) showing strong batch effects.

Supplemental FIGURE 4

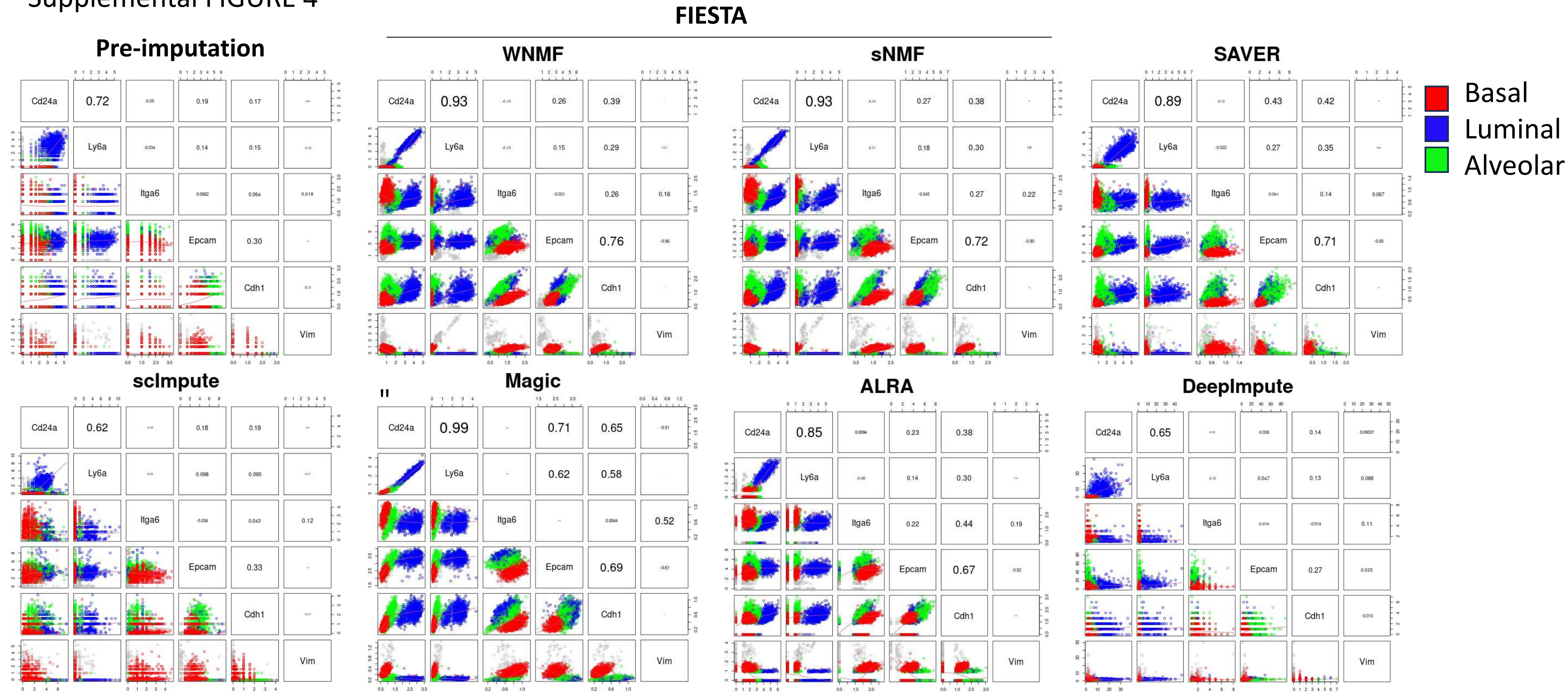


**Supplemental Figure 4.** Correlations (Pearson) and cell subtype specificity of select marker genes for adult mouse mammary cells (17) prior to imputation (upper left) and following imputation by various methods as indicated.

Supplemental FIGURE 5

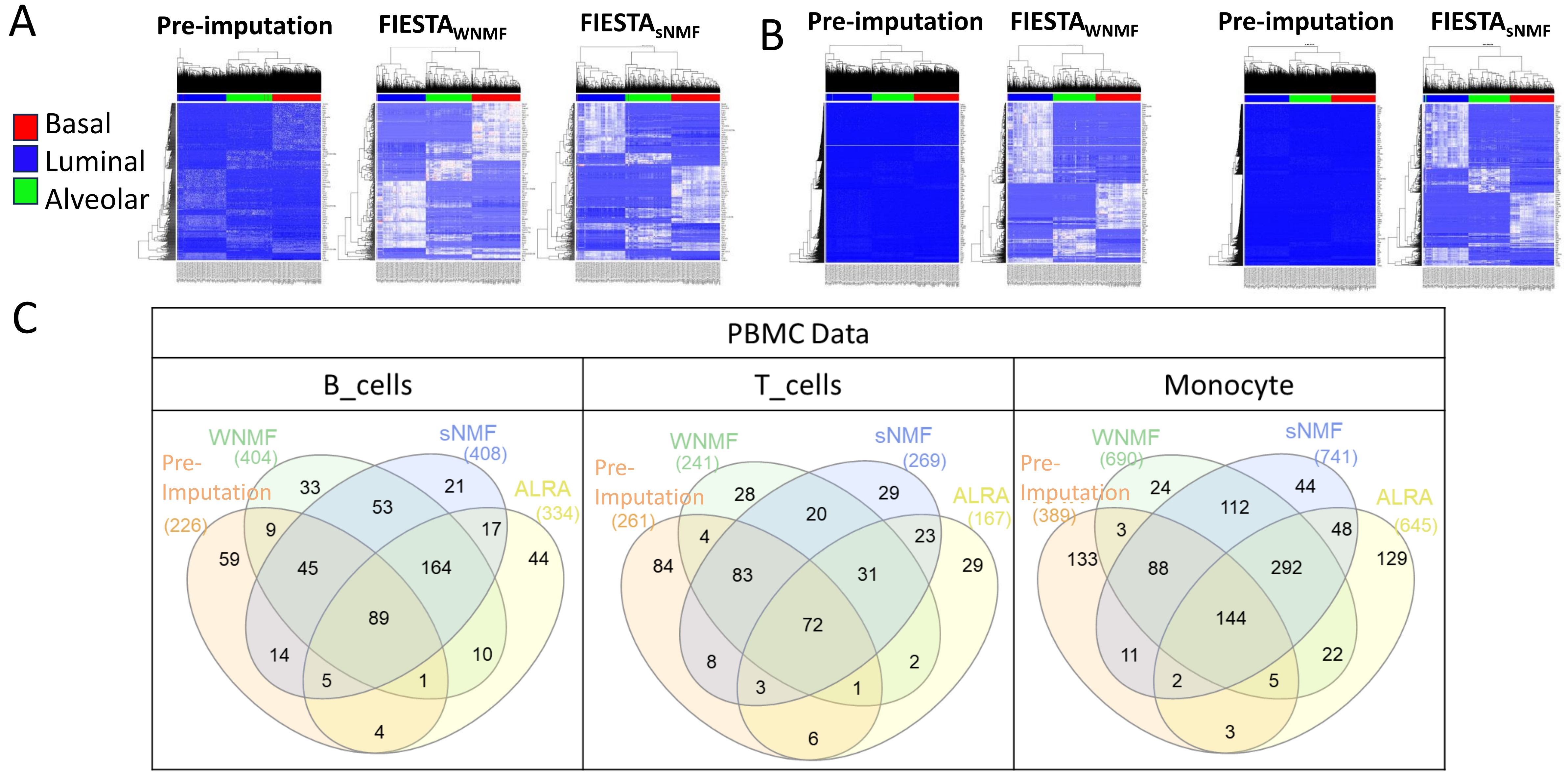


**Supplemental Figure 5.** A) Heat maps for DEG identified in pre-imputation adult mouse mammary data (17) vs. the same genes after FIESTA imputation. B) Genes identified as differentially expressed only after imputation with pre-imputation levels shown. C) The number and overlap of DEG identified prior to imputation or after FIESTA or ALRA.
